# Structural basis for the binding selectivity of human CDY chromodomains

**DOI:** 10.1101/820647

**Authors:** Cheng Dong, Yanli Liu, Tian-Jie Lyu, Serap Beldar, Kelsey N. Lamb, Wolfram Tempel, Yanjun Li, Zoey Li, Lindsey I. James, Su Qin, Yun Wang, Jinrong Min

## Abstract

The CDY (Chromodomain on the Y) family is a small family of chromodomain containing proteins, whose chromodomains closely resemble those in HP1 and Polycomb. The CDY proteins play an essential role in normal spermatogenesis and brain development. Dysregulation of their expression has been linked to male infertility and various neurological diseases. Like the chromodomains of HP1 and Polycomb, the CDY chromodomains also recognize the lysine-methylated ARKS motif embedded in histone and non-histone proteins. Interestingly, the CDY chromodomains exhibit different binding preferences for the lysine-methylated ARKS motif in different sequence contexts. Here, we present the structural basis for selective binding of CDY1 to H3K9me3 and preferential binding of CDYL2 to H3tK27me3 over H3K27me3. Based on our structural, binding and mutagenesis data, we synthesized a more CDYL1/2 selective peptidic ligand UNC4850. Our work provides critical implications that CDYL1b’s role in the regulation of neural development is dependent on its recognition of lysine-methylated ARKS motifs.

## Introduction

The *CDY* (Chromodomain on the Y) gene family originates from the autosomal gene *CDYL1* (CDY-like 1)^1,2^. Over the evolutionary timeline, the *CDYL1* gene was duplicated to create another autosomal gene *CDYL2*, and retroposed into the Y chromosome to generate *CDY*, which was further amplified into *CDY1* and *CDY2*. *CDY1* and *CDY2* are highly similar at both the DNA and protein sequence levels (>98% identity), and both genes contain two almost identical copies. The CDY family proteins contain two conserved domains: a canonical chromodomain in the N-terminus and a crotonase-like catalytic domain in the C-terminus (Fig. S1a and Fig. S2)^3^. The chromodomains of the CDY family are highly homologous to chromodomains in HP1 and Polycomb, and as such, they also recognize the lysine-methylated ARKS motif embedded in histone and non-histone proteins^4,5^. The catalytic domain of mouse CDYL1 has been reported to exhibit crotonyl-CoA hydratase activity^6^.

The CDY family of genes display diverse expression patterns. All human CDY family genes are abundantly expressed in testis (Fig. S1b), and their normal expression is required for complete spermatogenesis^7,8^. Deletion of either *CDY1* or *CDY2* leads to spermatogenic failure and male infertility^9–12^, and it has been proposed that *CDY2* and *CDY1* function in the early and later stages of spermatogenesis, respectively^7^. However, over the process of evolution, non-simian mammals have been shown to lack the Y-linked *CDY* genes^2^. Nevertheless, in mice, a short transcript of *CDYL1* displays testis-specific expression^1,2^, and the normal expression of mouse *CDYL1* is essential for spermatogenesis and male fertility, similar to human *CDY1/2* genes^6,13^. Both human and mouse CDYL1 function as transcriptional corepressors^14,15^. This CDYL1-mediated repressive function has recently been linked to its role in converting crotonyl-CoA, an active mark in transcription, to β-hydroxybutyryl-CoA^6^, and this crotonyl-CoA hydratase activity contributes to its role in spermatogenesis^6^. However, in another mouse study, conditional knockout of *CDYL1* does not affect global histone crotonylation, but causes disordered patterns of histone methylation and acetylation in testis^13^.

Mouse *CDYL1* is also involved in brain development, and its dysregulation has been linked to different neurological disorders^16–19^. For instance, CDYL1 regulates dendritic growth and morphology by repressing BDNF, a key regulator in dendrite development^16^; it modulates neuronal intrinsic plasticity by repressing SCN8A, a sodium channel gene whose mutations cause epileptic encephalopathies^17^; it controls neuronal migration by repressing RhoA, a gene involved in actin cytoskeleton regulation, and its deficiency causes neuronal migration disorders and increased susceptibility to epilepsy^18^; and it contributes to stress-induced depression by repressing VGF (or VGF nerve growth factor inducible), a gene that plays a key role in synaptic plasticity^19^. CDYL1 represses expression of its target genes through recruitment of PRC2-mediated H3K27me3 activity ^16,17,19^, and possibly also through its own histone crotonylation ability^19^. In addition to recruiting PRC2 and its associated H3K27me3 activity, CDYL1 also recruits G9a, a histone H3K9 dimethyltransferase, which together with REST represses transcription of potential tumor suppressor genes^15^. Not surprisingly, both CDYL and G9a are upregulated in hepatocellular carcinoma (HCC), but not in non-cancerous liver tissues^20^. CDYL1 also associates with chromatin assembly factor 1 (CAF1) and the MCM complex, and recruits histone H3K9/27 methyltransferases G9a, SETDB1, and PRC2 to the replication forks for transmission of repressive chromatin marks during DNA replication^21^, or to DNA double-strand break (DSB) sites for transcriptional silencing and promotion of homology-directed DNA repair (HDR)^22^.

The chromodomains of the CDY family bind to the lysine-methylated ARKS motif present in histones H3K9, H3K27 and H1.4K26 sites and in histone di-methyltransferase G9a *in vitro*^4,5^. Since CDYL1 has been shown to be associated with the lysine methyltransferases G9a, SETDB1 and PRC2, which are also able to methylate the lysine residue in the ARKS motif, it was proposed that CDYL1 and lysine methyltransferases, such as G9a and PRC2, synergize to propagate/transmit the H3K9me2 and H3K27me3 marks during DNA replication and transcriptional silencing^21,23,24^. Indeed, during X-chromosome inactivation, both H3K9me2 and H3K27me3 are required to recruit CDYL1 and G9a to the X chromosome, which together spread the repressive H3K9me2 marks along the X-chromosome^23^. Although we know that the chromodomains of the CDY family bind to both H3K9me3 and H3K27me3 (including both canonical histone H3K27me3 and testis-specific H3tK27me3), and both previous and our binding data presented here show that the chromodomain of CDY1 selectively recognizes H3K9me3 and the chromodomains of CDYL1 and CDYL2 preferentially bind to H3K9me3 and testis-specific H3tK27me3, the molecular mechanism underlying the binding selectivity of these chromodomains remains unclear. The present study elucidates the structural basis for the binding selectivity of the CDY chromodomains using histone peptides and a previously reported peptide-like inhibitor UNC3866^25^, as well as a more CDYL1/2 selective ligand UNC4850. Our results revealed that CDYL1 regulates neural development dependent on its readout of lysine-methylated ARKS motifs.

## Results and Discussion

### Chromodomains of the CDY family exhibit different binding selectivity to lysine methylated ARKS motif in different sequence contexts

The human genome contains six *CDY* genes: *CDY1A/B, CDY2A/B* and *CDYL1/2*. Among these *CDY* genes, the protein sequences of the chromodomains of the four Y-chromosome CDY proteins are identical (referred to as CDY1 later on for simplicity). The autosomal CDYL1 has three isoforms, CDYL1a, CDYL1b and CDYL1c (Fig. S1a), but only the most abundant CDYL1b harbors a methyllysine binding chromodomain and is recruited to the X chromosome coated with H3K9me2 and H3K27me3 marks, contributing to transcriptional repression^5,20,23^. As a result, we largely focused on CDYL1b in this study. The histone binding ability of these three unique and functional chromodomains (CDY1, CDYL1b and CDYL2) within the CDY family has been characterized previously by fluorescence polarization^4,5^. In this study, we further examined the binding affinities of these three chromodomains using the same length of peptides of H3K9me3, canonical H3K27me3, and testis-specific H3tK27me3 by ITC (Isothermal Titration Calorimetry) (Fig. 1a and b). We included the testis-specific H3tK27me3 in this study due to the important role played by the CDY proteins in spermatogenesis. During spermatogenesis, the haploid genome exhibits enhanced histone H3K9 and H3K27 methylation^26^. H3 contains at least five variants, including a testis-specific variant, H3t or H3.4^27^. The testis-specific H3t tail also contains an ARKS motif but carries a valine instead of a canonical alanine at position 24 (Fig. 1a).

**Figure 1.**
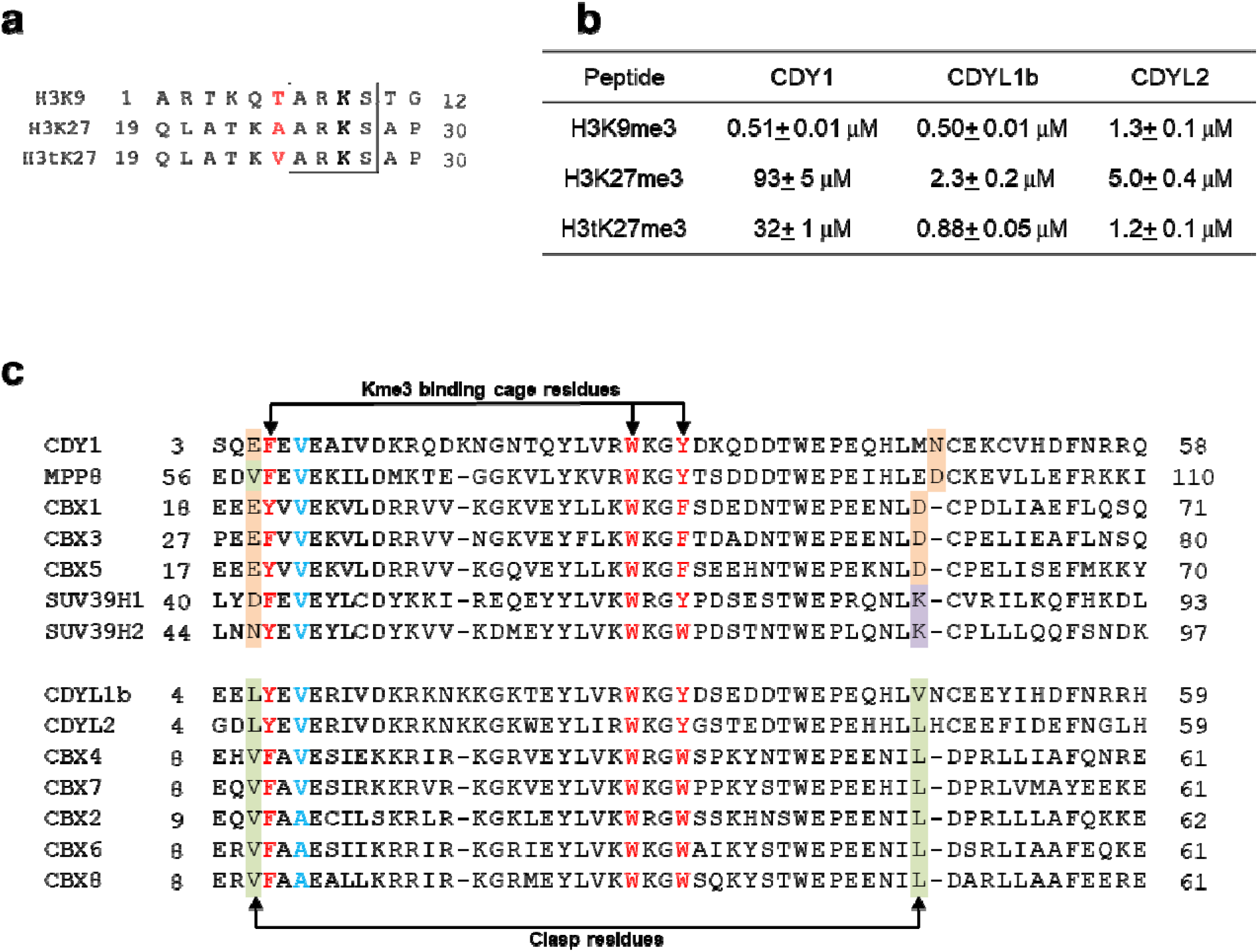
Chromodomains of the CDY family selectively bind to lysine methylated ARKS motif in different sequence contexts. **a**, Sequence comparison of ARKS motif-containing histone peptides. H3t, testis-specific histone H3. **b**, Binding affinities of selected CDY chromodomains to different lysine methylated ARKS motif-containing histone peptides determined by ITC. The *K_d_* errors were estimated from curve fitting of single measurements. **c**, Sequence alignment of selected chromodomains.

Our ITC binding results reveal that CDY1 is highly specific toward H3K9me3, with more than 60-fold selectivity over the H3tK27me3 and H3K27me3 peptides (*K*_*d*_ ~0.5 μM vs. 32 μM and 93 μM, Fig. 1b). On the other hand, CDYL1b and CDYL2 bind to these three peptides with a slight preference for H3K9me3 and H3tK27me3. Hence, the Y-chromosomal CDY1 is a specific reader of H3K9me3, whereas autosomal CDYL1b and CDYL2 are preferential binders of both H3K9me3 and testis-specific H3tK27me3 *in vitro*.

### Structural basis for selective binding of CDY1 to H3K9me3

To reveal why Y-linked CDY1 is a specific reader of H3K9me3, we determined the crystal structure of the CDY1 chromodomain in complex with an H3K9me3 peptide (Supplementary Table 1). Not surprisingly, the CDY1 chromodomain exhibits a canonical chromodomain fold, consisting of three β-strands followed by a C-terminal α helix, and binds to H3K9me3 in a similar manner to that observed in the complex structures of the HP1 and Pc subfamily of chromodomains with histone H3K9/K27 peptides^28–33^. The histone H3K9me3 peptide, together with the chromodomain’s N-terminus (β0), forms a short anti-parallel, double-stranded β-sheet on one side, and on the other side interacts with the chromodomain region just C-terminal of η2 (Fig. 2a). The ARKS motif is recognized through a conserved binding mode^28–33^. In brief, the K9me3 residue is recognized by a conserved aromatic cage formed by F6′, W28′ and Y31′ (Fig. 1c and 2b) (hereinafter, chromodomain residue numbers are marked with a prime (’) to distinguish them from the histone residue numbers). The histone H3A7 residue at the n-2 position (n represents the methyllysine residue) is buried in a hydrophobic pocket formed by V8′, V26′, W28′ and L44′ (Fig. 2b). The hydroxyl group of H3T6 forms a hydrogen bond with the carboxyl group of E5′ in CDY1. An equivalent hydrogen bond would be absent in binding H3K27me3 or H3tK27me3 which contain an alanine and valine at this position, respectively (Fig. 1a), which helps to explain the observed selectivity for the H3K9me3 sequence. CDY1 E5′ can additionally interact with the CDY1 N46′ side chain, located at the opposite edge of the peptide-binding groove (Fig. 2b). This arrangement to some extent resembles a pair of glutamic acid and aspartic acid residues in CBX1/3/5, which also exhibit strong selectivity toward H3K9me3 (Fig. 1c)^31^. Here, CDY1 also uses a similar E-N clasp for the selective H3K9me3 binding.

**Figure 2.**
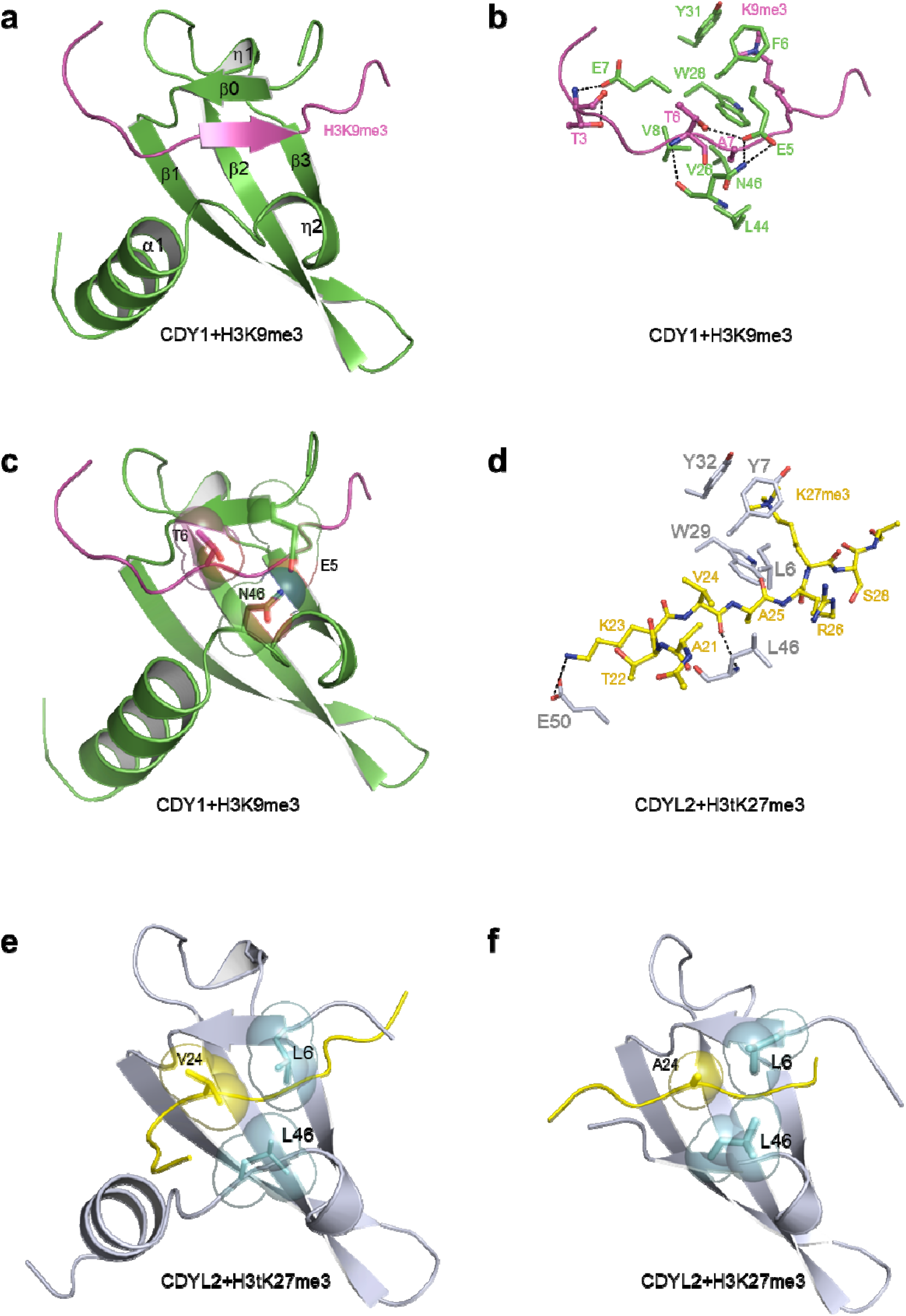
Structural basis for the selective binding of ARKS motif-containing histone peptides by CDY family chromodomains. **a-c**, Crystal structure of the CDY1-H3K9me3 complex. The peptide is colored in magenta and the protein is colored in green. **d-e**, Crystal structure of the CDYL2-H3tK27me3 complex. The peptide is colored in yellow and the protein is colored in gray. **f**, Crystal structure of the CDYL2-H3K27me3 complex, the C-terminal helix of CDYL2 is not shown due to its disorder caused by the artificial crystal packing. The interaction residues are shown in stick or sphere mode and the hydrogen bonds are shown as dashed lines.

### Structural basis for preferential binding of H3tK27me3 over H3K27me3 by CDYL2

Previous studies have extensively investigated the structural basis for the binding of the HP1 and Pc subfamily of chromodomains to the lysine-methylated ARKS motif in H3K9me3/H3K27me3^28–34^. However, it is still not clear how CDYL1b and CDYL2 achieve preferential binding to the testis-specific H3tK27me3. Because H3tK27me3 also harbors an ARKS motif, sequences immediately outside the ARKS motif might contribute to the binding preference. The H3tK27me3 peptide shows a single substitution, A24V, compared to the canonical H3K27me3 peptide (Fig. 1a). To see if the H3t-specific V24 immediately preceding the ARKS motif contributes to the binding preference, we determined the crystal structures of the CDYL2 chromodomain bound to the H3tK27me3 peptide or the H3K27me3 peptide, respectively (Supplementary Table 1, Fig. 2d-2f and Fig. S3a). The CDYL2 chromodomain in the CDYL2-H3tK27me3 complex displays a canonical chromodomain fold. However, in the CDYL2-H3K27me3 complex, the C-terminal α-helix of CDYL2 in one monomer is swayed away from its chromodomain core and occupies a similar position on another, crystal symmetry-related chromodomain molecule, thus forming a loose chromodomain homodimer (Fig. S3a). This unusual reorientation of the C-terminal α-helix and consequent dimerization may be caused by crystal packing, as our NMR solution structure of this chromodomain indicates a canonical chromodomain fold (Fig. S3b).

In the CDYL2-H3tK27me3 structure, V24 of the H3tK27me3 peptide packs against a “hydrophobic clasp” formed by L6′ and L46′ (referred to as L-L clasp as in reference^31^). V24 and R26 of the H3t peptide clasp the “L-L clasp” in turn (Fig. 2d). The methyl group of H3tA21 also packs against the H3tV24 side chain such that the H3t-specific residue V24 is largely buried in a hydrophobic groove (Fig. 2d and e). The equivalent methyl side chain of A24 in the canonical H3K27me3 peptide engages in less extensive hydrophobic interactions than the isopropyl side chain of H3tV24 (Fig. 2f). In the context of H3K9me3, the residue corresponding to V24 of H3tK27me3 is T6 (Fig. 2c). Valine and threonine differ by mere substitution of a hydroxyl for a methyl group in the amino acid’s γ position; a lack of strong selectivity between H3K9me3 and H3tK27me3 is thus not surprising.

### CDYL1b regulates neural development dependent on its recognition of lysine-methylated ARKS motifs

Our previous studies have shown that CDYL1b plays a vital role in neural development, and the gene repression of BDNF and RhoA mediated by CDYL1b is required for dendritic branching and neuronal migration^16,18^. Since the chromatin association of CDYL1b depends on its recognition of H3K9me2, H3K9me3 and H3K27me3 marks^5,16,21,23,24^, presumably a fully functional chromodomain is critical for CDYL1b to function properly in neural development. The use of a cell immunofluorescence staining assay revealed that flag-tagged CDYL1b colocalized with the H3K9me3 mark in the neuro-2a cell line (Fig. 3a and b). Our structural data show that the methyllysine-binding aromatic cage is composed of Y7′, W29′ and Y32′ in CDYL2 and is highly conserved in the CDYL1b (Fig. 1c and 2d). It is well known that the aromatic cage is essential in recognizing the lysine methylation marks^35^. Accordingly, when we mutated the cage residues of the CDYL1b chromodomain, such as Y7A′, W29A′ and Y32A′, CDYL1b is mislocalized and becomes diffuse in the nucleus (Fig. 3a-c), presumably due to the loss of its interaction with H3K9me3 (Fig. 3d). In contrast, mutations of residues that are not critical for binding including K30A′ or G31A′ did not affect the colocalization of CDYL1b and H3K9me3 (Fig. 3d). Hence, CDYL1b is localized to chromatin in the neuro-2a cells via the recognition of the lysine-methylated ARKS motif-containing histone marks.

**Figure 3.**
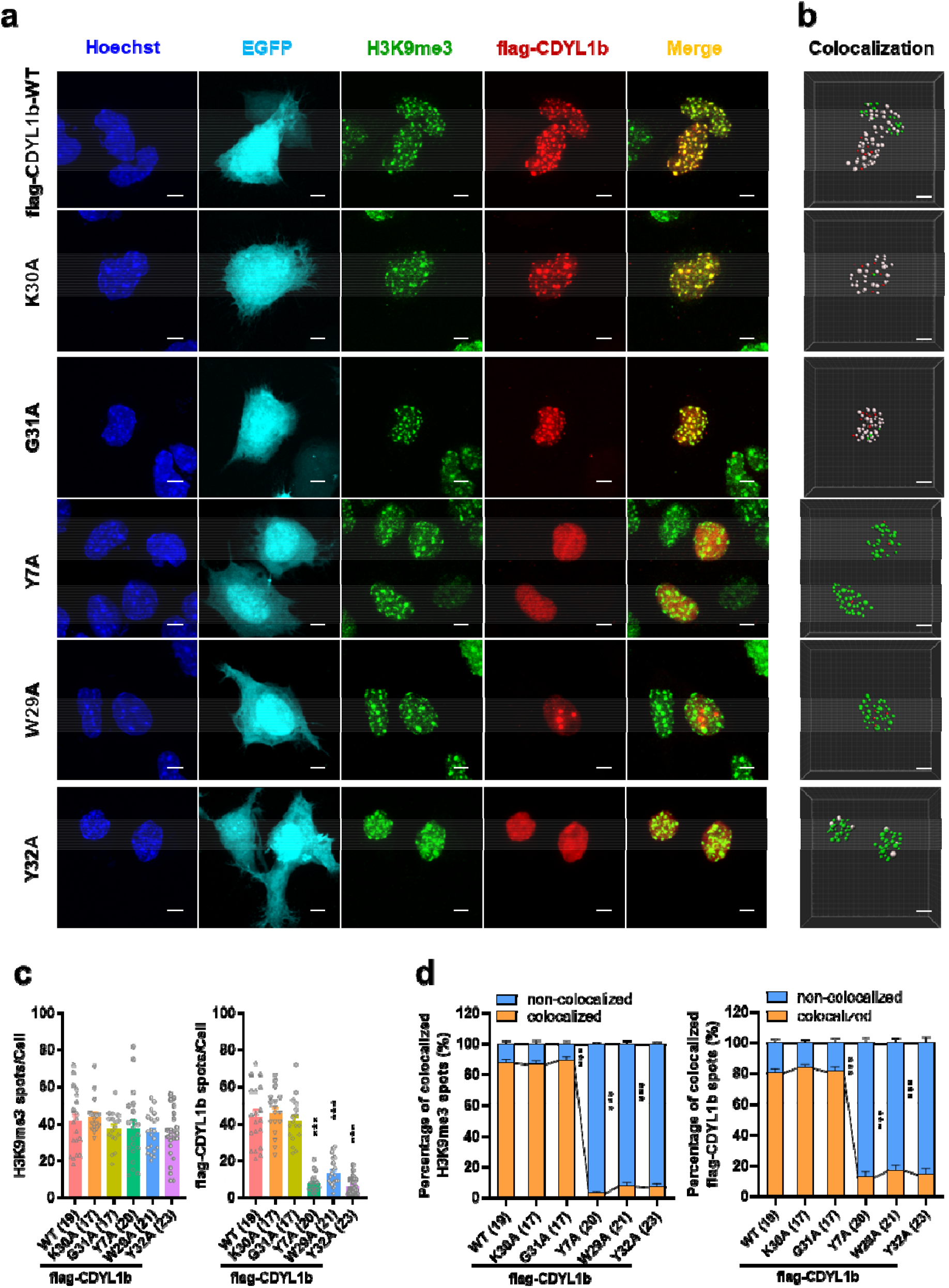
CDYL1b is associated with the heterochromatin mark H3K9me3 in the neuro-2a cell line. **a**, Representative confocal immunofluorescence images of EGFP positive cells stained with Hoechst, H3K9me3 antibody and flag antibody. Maximum intensity projections of EGFP (indigo, cell contour), Hoechst (blue, nucleus), H3K9me3 (green, heterochromatin), flag (red, flag-CDYL1b) and an overlay of H3K9me3 channel with flag channel are shown. Scale bar, 5 μm. **b**, Representative three-dimensional (3D) rendering of z-stack images from confocal microscope in (a) by using Imaris. The spots colocalization function of Imaris was used to split the spots into colocalized/non-colocalized spots. Green for non-colocalized H3K9me3, red for non-colocalized flag-CDYL1b and white for colocalized spots. Scale bar, 5 μm. **c**, Quantification of the number of H3K9me3 and flag-CDYL1b fluorescent spots per cells in (b). ***p<0.001, one-way ANOVA with Tukey’s multiple-comparisons test. At least 15 images per group were analyzed in two independent experiments. Data are presented as mean ± SEM. **d**, Spots colocalization analysis of images in (b), which shows summary of the percentage of colocalized H3K9me3 and flag-CDYL1b spots. ***p<0.001, two-way ANOVA with Tukey’s multiple-comparisons test. At least 15 images per group were analyzed in two independent experiments. Data are presented as mean ± SEM.

CDYL1b has also been shown to suppress dendritic morphogenesis through its association with chromatin^16^. Indeed, wild-type CDYL1b distinctly restricted dendritic branching, but its aromatic cage mutants Y7A′, W29A′ or Y32A′ were defective in the regulation of dendritic branching (Fig. 4a and b). As a control, the K30A′ mutant resembled wild-type CDYL1b. We next investigated whether CDYL1b regulates neuronal migration via binding to methylated histone marks. Knockdown of CDYL1b by a short hairpin RNA (shRNA) has been shown to result in defective neuronal migration^18^. We co-expressed CDYL1b-R-WT that is resistant to CDYL1b shRNA and showed that this is sufficient to restore neuronal migration. However, the Y7A′, W29A′ or Y32A′ mutants of CDYL1b-R failed to restore neuronal migration (Fig. 4c and d). Taken together, these results highlight the important role of CDYL1b in regulating neural development through its interaction with histone marks (H3K9me3/H3K27me3) by its functional chromodomain.

**Figure 4.**
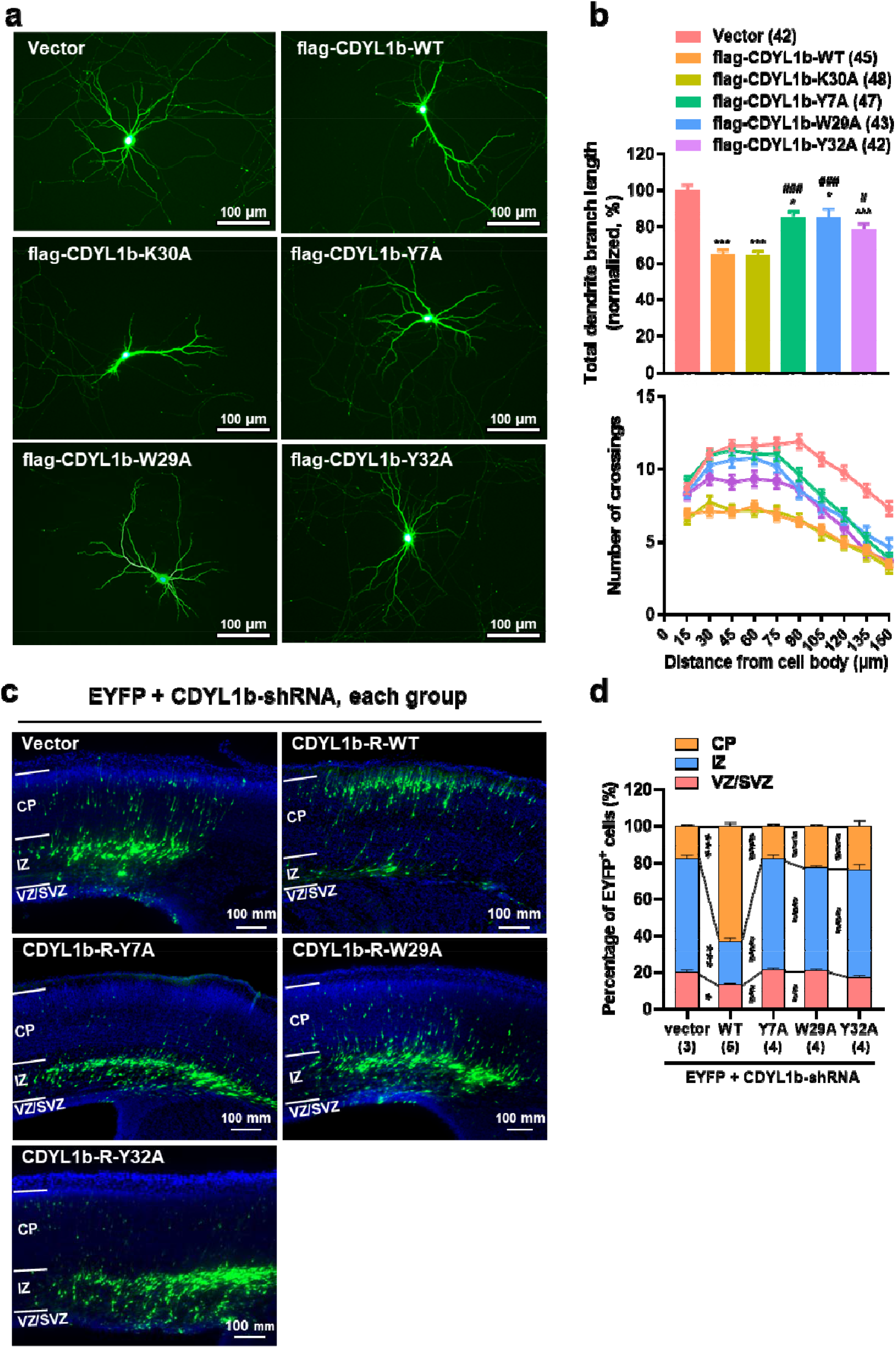
CDYL1b regulates neurodevelopment dependent on its readout of lysine methylated ARKS motifs. **a, b**, Negative regulation of dendritic branching by CDYL1b is dependent on the interaction between chromodomain of CDYL1b and lysine-methylated ARKS motifs. (a) Representative images of primary hippocampal neurons transiently transfected at DIV8 for 3 days with indicated plasmids. Neuronal morphology was visualized by cotransfection with pEGFP-N1. Scale bar, 100 μm. (b) Quantification of the total dendritic length (upper panel) and Sholl analysis of dendritic branching complexity (lower panel) of the neurons as shown in (a). *p<0.05, ***p<0.001 vs vector group and #p<0.05, ###p<0.001 vs CDYL1b-WT group, one-way ANOVA with Tukey’s multiple-comparisons test. At least 10 images per group per experiment were analyzed in three independent experiments. Data are presented as mean ± SEM. **c, d**, CDYL1b regulates neuronal migration dependent on the recognition of lysine-methylated ARKS motifs by the chromodomain. (c) Representative slices of E18.5 brains in utero electroporated of CDYL1b-shRNA at E14.5 together with blank vector, wild-type CDYL1b-resistant (CDYL1b-R-WT) or the aromatic cage mutant CDYL1b-resistant (CDYL1b-R-Y7A, CDYL1b-R-W29A or CDYL1b-R-Y32A). The cells were co-transfected with EYFP to visualize the distribution of transfected neurons (green), and cell nuclei were stained with DAPI (blue). Scale bar, 100 mm. (d) Quantification of the distribution of EYFP-positive neurons in VZ/SVZ, IZ and CP zones of brain slices as shown in (c). *p<0.05, ***p<0.001 vs vector group and ##p<0.01, ###p<0.001 vs WT (CDYL1b-R-WT) group, two-way ANOVA with Tukey’s multiple-comparisons test. More than 1, 000 neurons from three to five brains were analyzed in each group. Data are presented as mean ± SEM.

### Utilization of a CBX4/7-specific chemical probe UNC3866 to investigate the binding selectivity of chromodomains

UNC3866 is a peptidomimetic chemical probe which targets the Polycomb CBX family of chromodomains and binds to CBX4 and CBX7 most potently with a *K*_d_ of ~ 100 nM for both proteins. It was designed based on the H3K27me3 peptide and a SETDB1 fragment (GFALKme3S)^25,31,36,37^. Both previous^25,38^ and our current binding studies have shown that UNC3866 also binds to the chromodomains of the CDY subfamily and MPP8 with a preference for the CDYL1/2 chromodomains. In order to understand the binding specificity of UNC3866 by the ARKme3S binding chromodomains, we crystallized UNC3866 in complex with the chromodomains of CDYL2 and MPP8, in addition to the CBX2/4/7/8-UNC3866 structures we solved previously^25^. By analyzing all of the available structures of the ARKme3S binding chromodomains in complex with UNC3866, we found that the nature of the chromodomain clasp influences UNC3866 binding selectivity (Fig. 5). The hydrophobic clasps from the CDYL1/2 and CBX2/4/7/8 chromodomains make hydrophobic interactions with the Phe and Leu side chains of UNC3866 (Fig. 5a). In contrast, the CBX1/3/5 and SUV39H1/2 chromodomains have charged clasp pairs and could not form such hydrophobic interactions (Fig. 1c). Accordingly, the CBX1/3/5 and SUV39H1/2 chromodomains displayed negligible binding to UNC3866 (Fig. 5b). MPP8 has a mixed V-D clasp, which exhibits weaker hydrophobic interactions with UNC3866 based on its complex structure (Fig. 5a) and binds to UNC3866 with a significantly reduced affinity (Fig. 5b).

**Figure 5.**
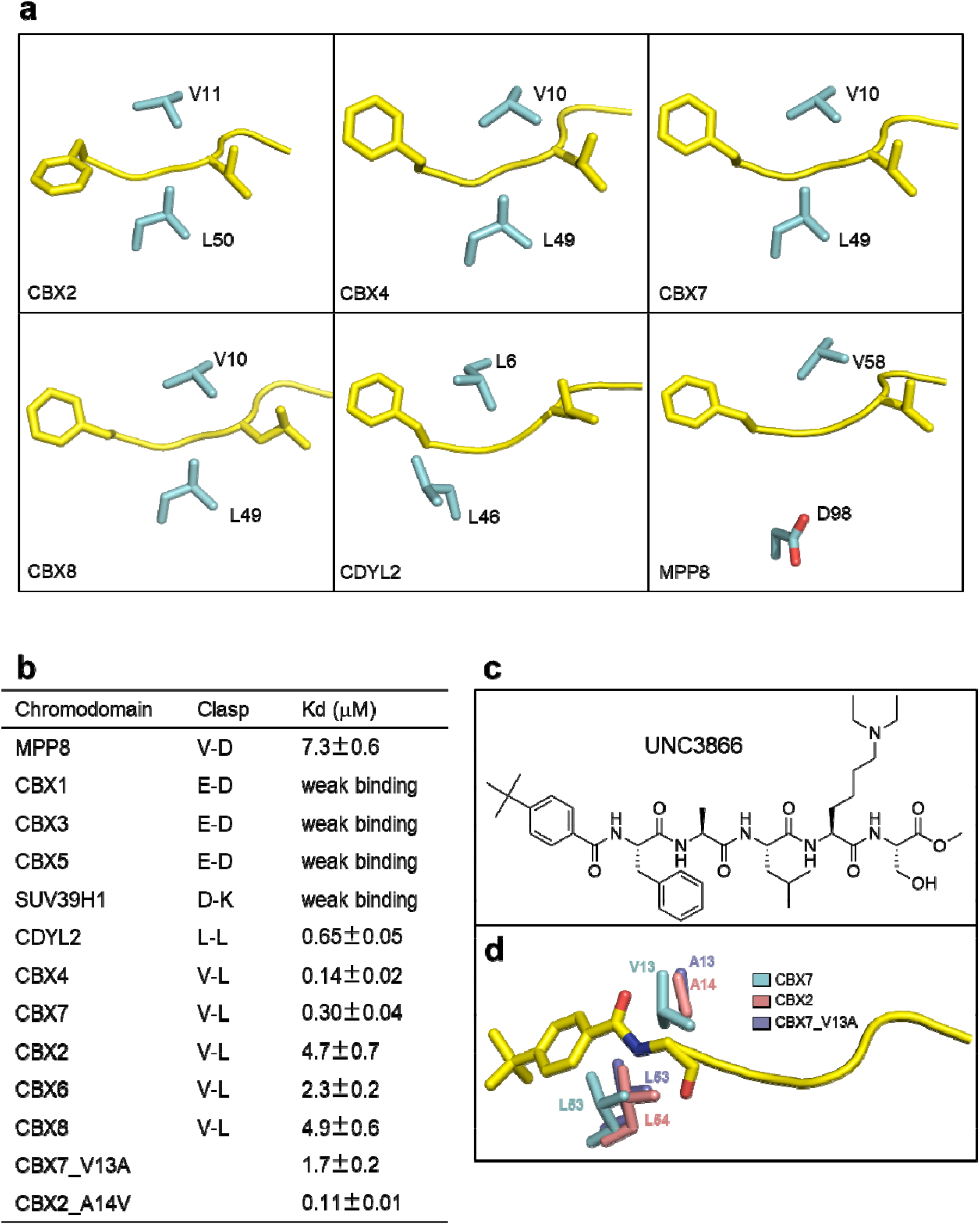
Investigation of binding selectivity of different chromodomains by using a CBX7 chemical probe UNC3866. **a**, Chromodomain clasp specific interactions from some representative chromodomains. **b**, Binding affinities of selected chromodomains to UNC3866 determined by ITC. The *K_d_* errors were estimated from curve fitting of single measurements. **c**, Chemical structure of UNC3866. **d**, Overlay of the hydrophobic groove residues of wild-type CBX2/7 and V13A′ CBX7 that recognize the tert-butylphenyl group of UNC3866.

In addition, the N-terminal tert-butylphenyl group of UNC3866 is accommodated in a largely hydrophobic groove in the CBX4/7 structures (Fig. 5d). Specifically, V13′ and L53′ of CBX7 packs snugly against the tert-butylphenyl group of UNC3866. But, in CBX2/6/8, an alanine residue replaces the V13′ residue of CBX7 (Fig. 1c), which may explain why CBX2/6/8 exhibit weaker binding to UNC3866. Like in CBX4/7, there is a valine at the corresponding position of the CDYL2 chromodomain (Fig. 1c), which may explain why CDYL2 shows higher binding affinity to UNC3866 than CBX2/6/8. In order to confirm this, we made the CBX7_V13A′ and CBX2_A14V′ mutants, and our binding results revealed that these substitutions reversed the binding affinities of CBX7 and CBX2 towards UNC3866 (Fig. 5b), with a greater than 40-fold increase in potency with CBX2_A14V′ over wild-type CBX2. This is further confirmed by our crystal structure of the CBX7_V13A′ mutant in complex with UNC3866. In this mutant structure, the alanine residue did not pack as snugly as the valine residue against the tert-butylphenyl group of UNC3866 (Fig. 5d). Taken together, this series of structures of UNC3866 in complex with different chromodomains has the potential to assist in the design of more selective chemical probes for these chromodomains in the future.

### Design and synthesize a CDYL1/2-specific compound UNC4850

Previously, we found that substituting the tert-butylphenyl capping group with an aliphatic isobutyl or cyclohexyl group resulted in increased selectivity for CDYL2 over CBX7, suggesting that chromodomain selectivity could be modulated at the N-terminal capping position^38^. And structural analysis reveals that, unlike in the CBX7^25^, H47′, E50′ and F51′ of CDYL2 form an incompact pocket packing against the tert-butylphenyl group of UNC3866, and hence, although the valine residue V9′ corresponding to V13′ of CBX7 is conserved, CDYL2 does not form strong interactions with the tert-butylphenyl group like CBX7 (Fig. 6a). Consequently, we synthesized and evaluated a similar ligand, UNC4850, which is a close analog of UNC3866 and contains an isobutyl N-terminal cap (Fig. 6b). As expected, UNC4850 binds to the chromodomain of CDYL1/2 about 10-fold more potently than to that of CBX7 by ITC (Fig. 6c). We also determined its complex structure with CDYL2. However, this complex structure displayed a similar crystal packing as we observed in the CDYL2-H3K27me3 complex (Fig. S3c), but unlike the UNC3866 complex structure. Notwithstanding this discrepancy, UNC3866 and UNC4850 occupy a conserved binding pocket (Fig. S3c). The smaller isobutyl cap of UNC4850 fits into the rearranged hydrophobic groove formed by V9′ and F51′ of CDYL2. In addition to a difference in the N-terminus, the leucine residue of UNC3866 is replaced with a phenylalanine residue in UNC4850, which is able to form stronger hydrophobic interactions with the conserved hydrophobic residues L6′ and L46′ (Fig. 6d). Therefore, UNC4850 is a specific small-molecule inhibitor toward CDYL1/2.

**Figure 6.**
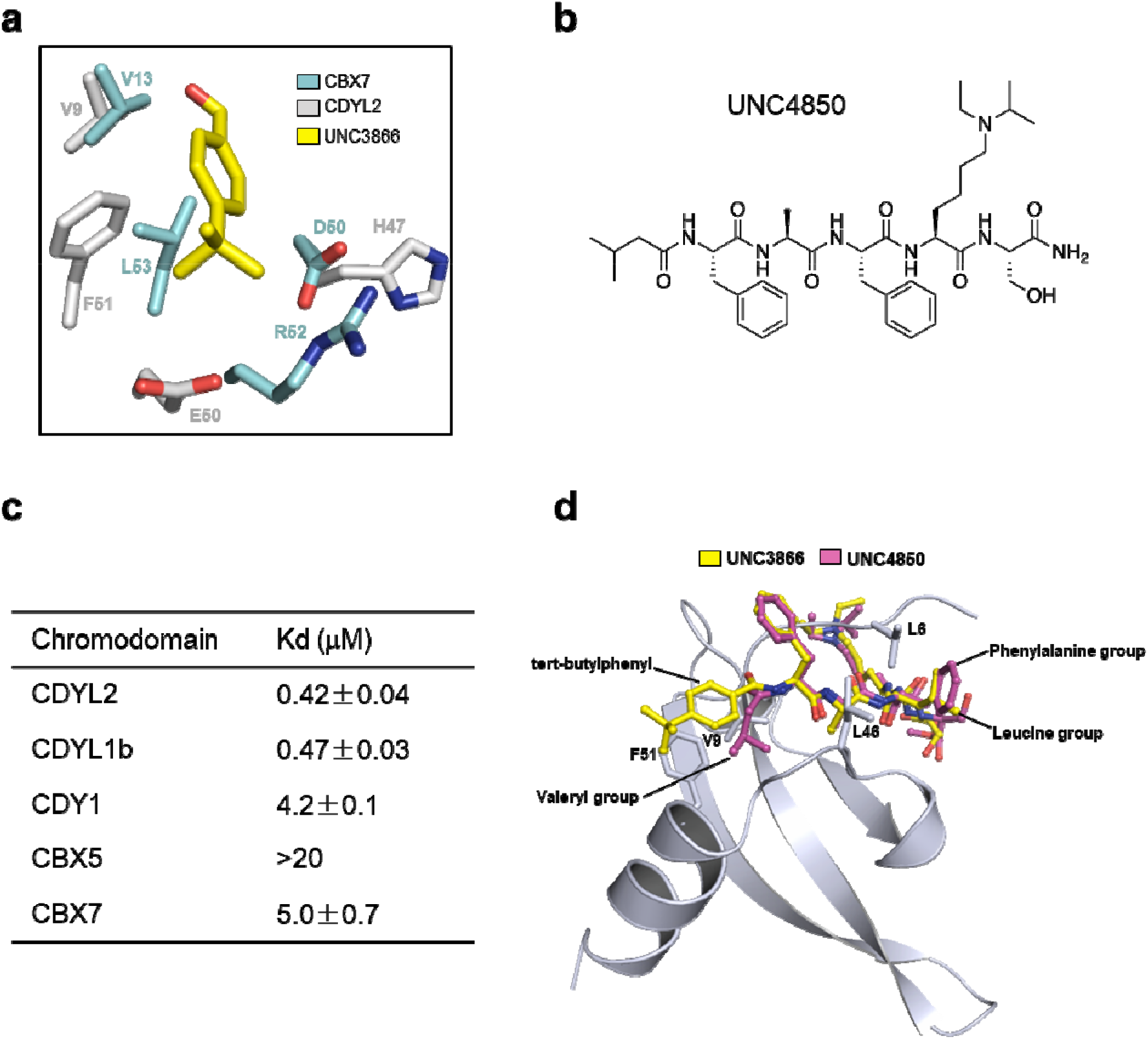
UNC4850 selectively recognizes CDYL1/2. **a**, Structural comparison of CDYL2 and CBX7 bound to the N-terminal tert-butylphenyl group of UNC3866. **b**, Chemical structure of UNC4850. **c**, Binding affinities of selected chromodomains to compound UNC480 determined by ITC. The *K*_*d*_ values were calculated as an average of at least two replicates ± standard deviation. **d**, Superimposition of CDYL2-UNC4850 complex and CDYL2-UNC3866 complex, only one CDYL2 molecule is shown for clarity.

## Methods

### Cloning, protein expression and purification

The following DNA fragments were cloned into a pET28a-MHL vector: CDY1 (aa 2-63), CDYL2 (aa 2-62 or 2-64), CBX2 (aa 9-62), CBX5 (aa 18-75), CBX7 (aa 8-62) and MPP8 (aa 56-116). CDYL1b (aa 1-66) was cloned into pET28GST-LIC. Protein expression was induced by 0.3 mM IPTG at 16 °C overnight, and proteins were purified by a Ni-NTA or GST column followed by ion exchange chromatography and size exclusion chromatography. His-tag or GST-tag was removed by the addition of TEV or Thrombin proteases. Uniformly ^15^N- and ^15^N/^13^C-labeled proteins were prepared by growing bacteria in M9 media using ^15^NH_4_Cl (0.5 g/liter) and ^13^C_6_-glucose (2.5 g/liter) as stable isotope sources. Mutations were introduced with the QuikChange II XL Site-Directed Mutagenesis Kit (Stratagene) and confirmed by DNA sequencing. Mutants were overexpressed and purified as the wild-type constructs above.

### Isothermal Titration Calorimetry

The ITC titrations between chromodomain proteins and UNC4850 were recorded at 25 °C with an Auto-iTC200 isothermal titration calorimeter (MicroCal Inc., USA). All protein and compound stock samples were prepared in an ITC Buffer (25 mM Tris-HCl, pH 7.5, 150 mM NaCl, and 2 mM β-mercaptoethanol) and then diluted to achieve the desired concentrations. Either 50 μM or 100 μM protein and 0.5 mM or 1.0 mM compound were used, respectively, to maintain a 10:1 compound to protein molar ratio for all ITC experiments. The concentration of the protein stock solution was established using the Edelhoch method, whereas compound stock solutions were prepared based on mass. A typical experiment included a single 0.2 μL compound injection into a 200 μL cell filled with protein, followed by 26 subsequent 1.5 μL injections of compound. Injections were performed with a spacing of 180 s and a reference power of 8 μcal/sec. The initial data point was routinely deleted. The other protein-ligands titrations were performed in a MicroCal VP-ITC instrument at 25 °C. The ITC buffer contains 20 mM Tris pH 7.5, 150 mM NaCl. The concentrations of protein samples and ligands range from 30-100 μM and 1-3 mM, respectively. The ligands were titrated into the protein solutions with 26 injections, containing 10 μL each spaced by 180 s with a reference power of 13 μcal/sec. The titration data was analyzed using Origin 7 Software (MicroCal Inc., USA) by nonlinear least-squares method, fitting the binding heats as a function of the compound to protein ratio in a one site-binding model.

### Protein crystallization

The chromodomains of CDYL2, CDY1, CBX7_V13A and MPP8 mixed with 1.5-3.0 fold peptide or compounds were crystallized using sitting drop vapor diffusion method by mixing 1 μL protein with 1 μL reservoir solution. The crystal of CDYL2 in complex with H3tK27me3 was obtained at 4 °C and the others were grown at 18°C. The CDYL2-H3K27me3 complex was crystallized in 1.6 M (NH_4_)_2_SO_4_, 10 mM MgCl_2_ and 0.1 M Sodium cacodylate trihydrate, pH 5.5; The CDYL2-H3tK27me3 complex was crystallized in 23% PEG3350, 0.25 M NH_4_H_2_PO_4_ and 0.1 M Sodium cacodylate trihydrate, pH6.0; The CDY1-H3K9me3 complex was crystallized in 1.4 M sodium citrate and 0.1 M Hepes, pH7.5; The CDYL2-UNC3866 complex was crystallized in 25% P3350, 0.2 M ammonium acetate and 0.1 M Hepes, pH 7.5; The CDYL2-UNC4850 complex was crystallized in 15% PEG8000, 0.2 M MgCl_2_ and 0.1 M Tris-HCl, pH8.5; The CBX7 (V13A)-UNC3866 complex was crystallized in 1.5 M (NH_4_)_2_SO_4_ and 0.1 M Bis-Tris propane, pH 7.0; The MPP8-UNC3866 complex was crystallized in 0.2 M MgCl_2_ and 0.1 M Tris-HCl, pH 8.5.

### Crystal diffraction data collection and structure determination

CDY1-H3K9me3: Diffraction data were collected on a copper rotating anode source. XDS^39^, AIMLESS^40^ and XIA2^41^ were used for data reduction. The structure was solved by molecular replacement with the program PHASER and coordinates from PDB entry [4HAE]. The model was automatically rebuilt with the program ARP/wARP^42^. COOT^43^ and REFMAC^44^ were used for model interactive rebuilding and restrained refinement, respectively.

CDYL2-H3tK27me3: Diffraction data were collected at beamline 08ID of the Canadian Lightsource^45^ and reduced alternatively with DENZO/SCALEPACK^46^, XDS or XIA2^39,41^. COOT was used for interactive model rebuilding. PHENIX and REFMAC were used for restrained model refinement^44,47^. An anomalous scatterer near the protein constructs N-terminus was tentatively assigned as a nickel cation. The geometry of that ion’s coordination was restrained based on the results of a MOGUL^48^ search.

CDYL2-H3K27me3: Diffraction data were collected at beamline 19ID of the Advanced Photon Source and reduced alternatively with DENZO/SCALEPACK or XDS. The structure was solved by molecular replacement with the program PHASER^49^ and coordinates from PDB entry 4HAE. ARP/wARP was used for map improvement^50^. COOT was used for interactive model building. REFMAC and AUTOBUSTER^51^ were used for restrained model refinement.

CDYL2-UNC3866: Diffraction data were collected on a copper rotating anode source. Inspection of the set of diffraction images revealed the emergence of ice rings in the course of data collection. Data were initially processed with DENZO/SCALEPACK, and later with the XIA2 XDS/AIMLESS pipeline applying the “exclude_ice_regions” option. Molecular replacement was performed with PHASER and an ensemble comprising of preliminary versions of PDB entries 5EPJ and 5EPK^25^. After phase modification with ARP/wARP and PARROT^52^, the model was traced with the BUCCANEER pipeline^53^. Further model rebuilding and refinement were performed with COOT and REFMAC, respectively. Geometry restraints for UNC3866 were prepared using GRADE^54^, MOGUL and JLIGAND^55^.

CBX7-(V13A)-UNC3866: Diffraction data were collected on a copper rotating anode source and processed with the XIA2 XDS/AIMLESS pipeline. Preliminary coordinates of isomorphous PDB entry 5EPJ were used as a starting model for restrained refinement with REFMAC and interactive rebuilding with COOT.

MPP8-UNC3866: Diffraction data were collected on a copper rotating anode source and processed through the XIA2 XDS/AIMLESS pipeline. The structure was solved by molecular replacement with the program PHASER and coordinates from PDB entry 3R93^38^. COOT^43^ and REFMAC^44^ were used for model interactive rebuilding and restrained refinement, respectively. CDYL2-UNC4850: Diffraction data were collected at APS beamline 22ID and reduced with DENZO/SCALEPACK or, for later stages of model refinement, the XIA2 XDS/AIMLESS pipeline. The structure was solved by molecular replacement with the program PHASER and a search model derived from the CDYL2-UNC3866 complex (see above). The model was rebuilt automatically with PHENIX AUTOBUILD. COOT, REFMAC and PHENIX were used for additional rebuilding and refinement of the model. Some geometry restraints for the UNC4850 ligand were prepared on the GRADE server (http://www.globalphasing.com).

### NMR structure determination

Solution structure of the CDYL2 chromodomain was determined using the ABACUS approach^56^ from NMR data collected at high resolution from nonlinearly sampled spectra and processed using multidimensional decomposition^57,58^. NMR spectra were recorded at 25 °C on Bruker Avance 800-MHz spectrometers equipped with cryoprobes. The sequence-specific assignment of ^13^C, ^1^H, and ^15^N resonances were assigned using the ABACUS protocol^56^ from peak lists derived from manually peak picked spectra. The restraints for backbone φ and ψ torsion angles were derived from chemical shifts of backbone atoms using TALOS+^59^. Hydrogen bond constraints were used within regular secondary structure elements. Automated NOE assignment^60^ and structure calculations were performed using CYANA (Version 3.0) according to its standard protocol. The final 20 lowest-energy structures were refined within CNS1.3^61^ by a short constrained molecular dynamics simulation in explicit solvent^62^.

### Plasmids for cellular assays and *in utero* electroporation

Flag-tagged human CDYL1b expression plasmid^16^ (flag-CDYL1b-WT) and shRNA-resistant CDYL1b in pCAGGS-IRES-EGFP vector^18^ (CDYL1b-R-WT) were used to create the negative control mutants (flag-CDYL1b-K30A and flag-CDYL1b-G31A) and the aromatic cage mutants of CDYL1b (flag-CDYL1b-Y7A, flag-CDYL1b-W29A, flag-CDYL1b-Y32A, CDYL1b-R-Y7A, CDYL1b-R-W29A and CDYL1b-R-Y32A) by the Hieff Mut™ Site-Directed Mutagenesis Kit (<u>11003ES10</u>, Yeasen Biotech, China). Short hairpin RNA (shRNA) plasmid specific for CDYL1b (CDYL1b-shRNA, target GAGATATTGTCGTCAGGAA), pEGFP-N1 and EYFP used in the experiments were the same as described previously^18^.

### Cell culture and transfection

Neuro-2a cell line was purchased from ATCC and maintained in Dulbecco’s modified Eagle’s medium (DMEM) (CM15019, M&C Gene Technology, China) supplemented with 10% fetal bovine serum (FBS) (SA212.02, CellMax, China) at 37 °C in a humidified atmosphere of 5% CO_2_. Transfection was done with Lipofectamine 2000 (11668027, Invitrogen, USA), according to the manufacturer’s instructions. Neuro-2a cells were seeded on coverslips and co-transfected pEGFP-N1 plasmid for 6 hours with wild-type flag-CDYL1b (WT), or the negative control mutant flag-CDYL1b (K30A or G31A), or the aromatic cage mutant flag-CDYL1b (Y7A, W29A or Y32A).

### Immunofluorescence Staining

Cells were washed with 0.1 M PBS twice and then fixed in 4% paraformaldehyde (PFA) for 10 min. After three times 5 min washing with 0.1 M PBS, cells were permeabilized with 0.1% Triton™ X-100 in 0.1 M PBS for 20 min at room temperature and then blocked with 3% bovine albumin (BSA) in 0.1 M PBS with 0.3% Triton™ X-100 for 1 hour at room temperature. Next, cells were incubated overnight at 4 °C with monoclonal anti-FLAG M2 antibody (1:1000, F1804, Sigma-Aldrich, Germany) and polyclonal rabbit anti-H3K9me3 antibody (1:100, A2360, ABclonal, China). Next day, after three times 5 min washing with 0.1 M PBS, cells were incubated for 1 hour at room temperature with donkey anti-mouse Alexa Fluor 594 antibody (1:1000, A-21203, Thermo Fisher Scientific, USA) and donkey anti-rabbit Alexa Fluor 555 antibody (1:1000, A-31572, Thermo Fisher Scientific, USA). Finally, cells were counterstained with Hoechst 33342 (10 μg/ml, C0031, Solarbio, China) and mounted with antifading mounting medium (S2100, Solarbio, China).

### Colocalization analysis

Cells were captured at a 63×/1.40 NA Silicon Immersion objective lens with 4× digital zoom (512×512, pixel size in XY = 90.3 nm, Z-step = 0.3 μm) by TCS SP8 (Leica, Germany) laser-scanning confocal microscope. Z-stacks of confocal images were uploaded into the Huygens Professional software (Scientific Volume Imaging, The Netherlands) for deconvolution to make sure the images are suitable for the colocalization analysis. Then red (flag-CDYLb) and green (H3K9me3) channels were used to generate the three-dimensional (3D) spots by the Imaris ×64 9.0.1 software (Bitplane, Switzerland)^63^. First, Surface function of Imaris was used on the red (flag-CDYL1b) channel to create the regions of interest (ROIs) automatically. Next, spots function of Imaris was used on the red (flag-CDYL1b) and green (H3K9me3) channels to produce heterochromatin foci spots in automatic mode and followed by the “Split Into Surface Objects” and “Colocalize spots” XTensions of Imaris to measure the number of spots per cell and colocalized/non-colocalized spots in each channel. These data were used to estimate spots distribution pattern and make colocalization analysis. At least 15 images per group were analyzed in two independent experiments.

### Primary neuron culture, transfection and reagents

Primary mouse hippocampal neurons were prepared as previously described^16^. E16.5 ICR-strain pregnant mice checked for vaginal plugs were provided by the Animal Center of the Peking University Health Science Center. Hippocampal explants were isolated from E16.5 ICR-strain embryos of either sex at 4 °C (soaking in cold Hank’s Balanced Salt Solution on an open platform ice-free cooler XT Starter (BCS-504, BioCision, USA)). Tissues were then digested with 1 mL 0.25% trypsin in a 35 mm culture dish for 30 min at 37 °C with gently shaking the culture dish every 5 min. Then digested tissues were triturated with a pipette in plating medium (DMEM with 10% FBS and 100 μg/mL antimicrobial agent Primocin™ (ant-pm-1, Invivogen, USA). Dissociated neurons were immediately plated onto 35 mm dishes coated with 50 μg/mL poly-D-lysine (P6407, Sigma Aldrich, Germany), at a density of 5×10^5^ live cells with 2 mL plating medium per dish. About 4 hours later, the plating medium was changed to neurobasal medium (21103049, Gibco™, USA) supplemented with 2% B27 (17504044, Gibco™, USA) and 0.5 mM GlutaMAX-I (35050061, Gibco™, USA). Neurons were cultured at 37°C in a humidified atmosphere of 5% CO_2_ and fed by aspirating half of the medium from each well and replacing it with fresh complete Neurobasal medium every third day. To estimate the role of CDYL1b aromatic cage residue mutants in dendritic branching, neurons were co-transfected with the DNA of interest and pEGFP-N1 at day 8 *in vitro* (DIV8) for 20 min by Lipofectamine 2000, according to the manufacturer’s instructions.

### Neuronal morphology analysis

Neuronal morphology analysis was performed as previously described^16,64^. Prepared neurons were captured at a 20×magnification objective lens by a DMI4000 (Leica, Germany) inverted fluorescence microscope. Dissociated neurons grown at low density were used to determine morphological characteristics. Neurons transfected with DNA were selected by EGFP expression. The fluorescence images captured by DMI4000 were unloaded into Imaris for Sholl analysis^65,66^ of neuronal morphology. Filaments function of Imaris was used to trace and measure all the dendritic branches of EGFP positive neurons with AutoPath mode semi-automatically. Then Sholl analysis was performed on the data exported by Imaris, and the concentric circles (Sholl spheres Resolution) was defined as 15 μm differences in diameter. The total dendritic length and number of dendrites crossing each circle were measured to estimate dendritic branching complexity. The experimental group was normalized to the control group, which was set to 100% for convenience. At least 10 images per group per experiment were analyzed in three independent experiments.

### In utero electroporation

In utero electroporation was performed on E14.5 mouse brains as previously described^18^. E14.5 ICR-strain pregnant mice checked for vaginal plugs were provided by the Animal Center of the Peking University Health Science Center. The pregnant mice were anesthetized by intraperitoneal injection with 0.7% sodium pentobarbital (100 mg/kg). Mouse embryos were exposed in the uterus and 1 μL of DNA solution with 0.01% Fast Green was injected into the lateral ventricles of embryos through a sharped micropipette by a microelectrode beveler BV-10 (Sutter Instrument, USA). Plasmids were prepared in 0.9% sterilizing saline at the concentrations of 1 μg/μL for EYFP, 3 μg/μL for CDYL1b-shRNA, and 6 μg/μL for blank vector, wild-type CDYL1b-resistant (CDYL1b-R-WT) or the aromatic cage mutant CDYL1b-resistant (CDYL1b-R-Y7A, CDYL1b-R-W29A or CDYL1b-R-Y32A). Electrical pulses of 36 V were generated with an Electro Squire portator T830 (BTX, USA) and applied to the cerebral wall for 50 ms each, for a total of five pulses, at intervals of 950 ms. The uterus was then replaced, and the abdomen wall and skin were sutured. After surgical manipulation, pregnant mice were allowed to recover to consciousness in a 37 °C incubator.

Four days after surgery, E18.5 mice were perfused with 0.1 M PBS followed by 4% PFA and brains were fixed overnight in 4% PFA at 4 °C. Fixed brains were dehydrated and cryoprotected in 20% follower by 30% sucrose both for 24 hours, embedded in Tissue-Tek O.C.T. compound (<u>4583</u>, SAKURA, USA) in a −80 °C refrigerator, and then sectioned with a Leica CM3050 S Research Cryostat (Leica, Germany). Prepared brain sections were washed three times with 0.1 M PBS and mounted by antifading mounting medium (with DAPI). The mounted brain slices were scanned by a Virtual Slide Microscope VS120 (Olympus, Japan) with a 20× magnification objective lens. The images were uploaded into Imaris software to estimate the effect of DNA transferred by electroporation on neuronal migration. First, Surface function of Imaris was used on the blue (DAPI) channel to create the regions of interest (ROIs) in manual mode to split VZ/SVZ, IZ and CP zones of brain. Next, spots function of Imaris was used on the green (EYFP) channel and followed by the “Split into Surface Objects” XTensions of Imaris to measure the distribution of EYFP positive neurons in deferent zones. More than 1, 000 neurons from three to five brains were analyzed in each group.

### Statistical analysis

Data were analyzed and plotted using GraphPad Prism 6.0 (GraphPad Software, USA). Statistical methods for comparisons were chosen properly from unpaired t-test and one-way or two-way ANOVA with Tukey’s or Bonferroni’s multiple-comparisons test as described in figure legends. The statistical credibility was defined as three level: *p<0.05, *p<0.01 and ***p<0.001. All data are presented as mean ± SEM.

### UNC4850 Synthesis and Purification

Fmoc-protected amino acids were purchased from Chem-Impex and Sigma-Aldrich with the exception of the Fmoc lysine derivative which was synthesized as described previously^38^. All other chemicals and solvents were purchased from TCI America and Sigma Aldrich, unless otherwise stated. Syringe reaction vessels were made in-house with 2 mL syringes (Norm-Ject) and frits (Teledyne).

Synthesis was conducted via solid-phase peptide synthesis on Fmoc Rink amide resin (100 mg, Chem-Impex). The resin was initially swollen in DCM followed by DMF (10 minutes each). Fmoc deprotection was conducted by incubation with a solution of 2.5% 1,8-diazabicycloundec-7-ene and 2.5% pyrrolidine in DMF for 10 min. The resin was filtered and washed twice with DMF, methanol, DMF, and DCM before amino acid for coupling. Standard Fmoc synthesis protocols were used to generate the 6-mer peptide, UNC4850. Briefly, Fmoc-protected amino acids (4 eq) were mixed for 5 minutes with HBTU (4 eq), HOAt (4 eq), and DIPEA (8 eq) in 1 mL of DMF and 1 mL of dichloromethane (DCM). The solution was then added to the resin and left on a shaker at room temperature for 1 hour. The resin was filtered and washed twice with DCM, DMF, methanol, and DMF again. Fmoc amino acid protecting groups were removed as described above, and then the resin was filtered and washed twice with DMF, methanol, DMF, and DCM before adding the next amino acid for coupling. Following installation of the isovaleric acid capping residue, the resin was rinsed 6 times with DCM. Cleavage cocktail (95% trifluoroacetic acid, 2.5% triisopropylsilane, and 2.5% water) was added to the resin, the mixture was left on the shaker for 2 hours, and the filtrate was collected. The resin was rinsed twice with DCM and the filtrates were pooled and concentrated under vacuum.

The crude material was purified by preparative HPLC using an Agilent Prep 1200 series with the UV detector set to 220 nm and 254 nm. Samples were injected onto a Phenomenex Luna 250 × 30 mm, 5 μm, C18 column at 25 °C. Mobile phases of A (H_2_O + 0.1% TFA) and B (CH_3_CN) were used with a flow rate of 40 mL/min. Product fractions were pooled and concentrated to yield 19.1 mg (51.3% yield) of UNC4850 as a white solid.

Analytical LCMS and ^1^H NMR were used to establish purity. Analytical LCMS data was acquired using an Agilent 6110 Series system with the UV detector set to 220 nm and 254 nm. Samples were injected (10 μL) onto an Agilent Eclipse Plus 4.6 × 50 mm, 1.8 μm, C18 column at 25 °C. Mobile phases A (H_2_O + 0.1% acetic acid) and B (CH_3_OH + 0.1% acetic acid) were used with a linear gradient from 10% to 100% B in 5.0 min, followed by a flush at 100% B for another 2.0 min at a flow rate of 1.0 mL/min. Mass spectra (MS) data were acquired in positive ion mode using an Agilent 6110 single quadrupole mass spectrometer with an electrospray ionization (ESI) source. Nuclear Magnetic Resonance (NMR) spectra were recorded on a Varian Mercury spectrometer at 400 MHz for proton (^1^H NMR); chemical shifts are reported in ppm (δ) relative to residual protons in deuterated solvent peaks. Due to intramolecular hydrogen-bonding, hydrogen-deuterium exchange between the amide protons of the molecule and the deuterated solvent is slow and requires overnight equilibration for complete exchange. The characterization of UNC4850 was shown in Fig S4.

## Acknowledgments

We thank Dr. John R. Walker for reviewing some of the crystal structures. The SGC is a registered charity (number 1097737) that receives funds from AbbVie, Bayer Pharma AG, Boehringer Ingelheim, Canada Foundation for Innovation, Eshelman Institute for Innovation, Genome Canada through Ontario Genomics Institute [OGI-055], Innovative Medicines Initiative (EU/EFPIA) [ULTRA-DD grant number 115766], Janssen, Merck KGaA, Darmstadt, Germany, MSD, Novartis Pharma AG, Ontario Ministry of Research, Innovation and Science (MRIS), Pfizer, São Paulo Research Foundation-FAPESP, Takeda, and Wellcome. Some diffraction experiments were performed at the Structural Biology Center and Northeastern Collaborative Access Team beam lines at the Advanced Photon Source at Argonne National Laboratory. ANL is operated by the University of Chicago Argonne, LLC, for the U.S. Department of Energy Office of Biological and Environmental Research under contract DE-AC02-06CH11357. This work was also supported by a NSERC grant RGPIN-2016-06300 (JM), and the National Natural Science Foundation of China grants (31500615 (YL), 31500613 (SQ), 31530028, 31720103908 and 81821092 to Y.W., 31900865 (C.D.)), and the Central China Normal University (CCNU) from the college’s basic research and operation of Ministry of Education (MOE) [grant number CCNU18TS020]. This work was supported by the National Institute on Drug Abuse, National Institutes of Health (NIH) (Grant R61DA047023-01) and the University Cancer Research Fund, University of North Carolina at Chapel Hill to L.I.J. We would like to acknowledge the assistance of the microscope (Leica TCS SP8 and Olympus VS120) and software (Imaris 9.0.1, Bitplane), applied by Medical and Healthy Analytical Center of Peking University, the Imaging Facility of National Institute of Biological Sciences and the Imaging Core Facility, Technology Center for Protein Sciences of Tsinghua University in Beijing, China.

## Author contributions

C.D., Y.L. and S.Q. purified and crystallized the proteins and conducted the ITC assays; S.Q. performed the NMR experiment; T.L. and Z.L. performed the cellular and *in vivo* assays; W.T. and C.D. collected diffraction data and determined the crystal structures; K.N.L. synthesized the compounds and conducted ITC; Y.L. cloned the constructs; L.I.J, Y.W. and J.M. supervised the study; J.M. conceived the study and S.Q. and J.M. wrote the paper with substantial contributions from all the other authors.

