## Supplemental information for "Structural basis for the binding selectivity of human CDY chromodomains"

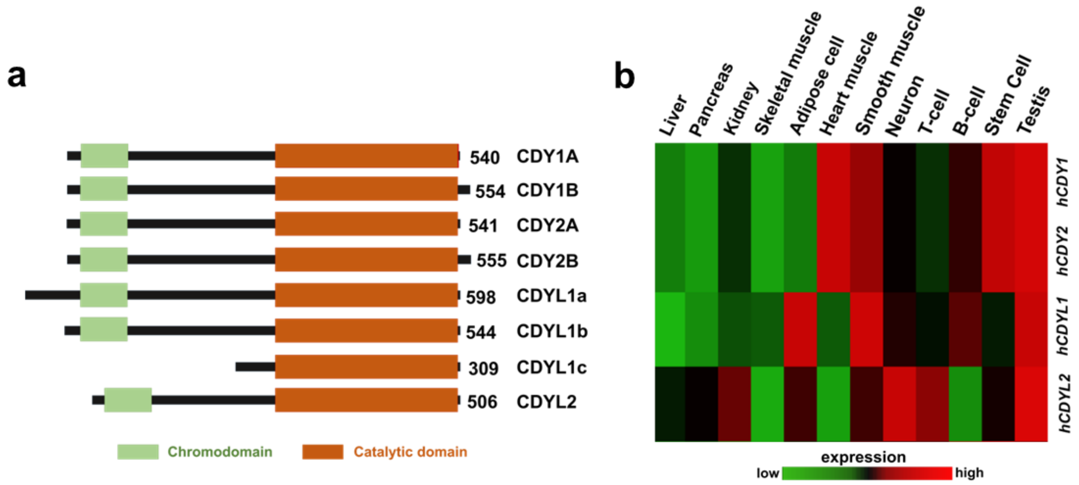


**Supplementary Figure 1** a, Domain architecture of human CDY family proteins. b, Tissue distribution of human CDY family genes.

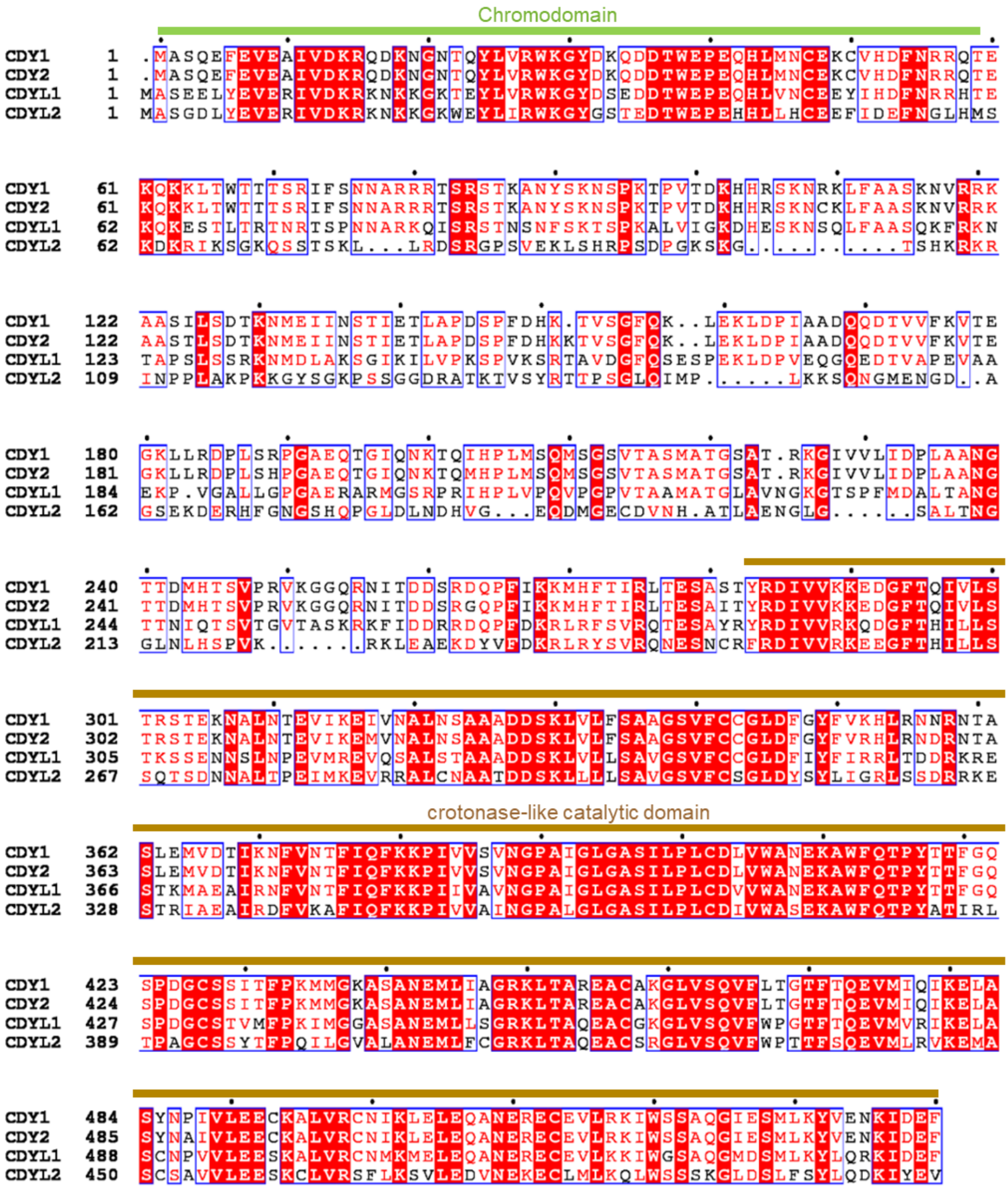


**Supplementary Figure 2** Sequence alignment of selected CDY family proteins

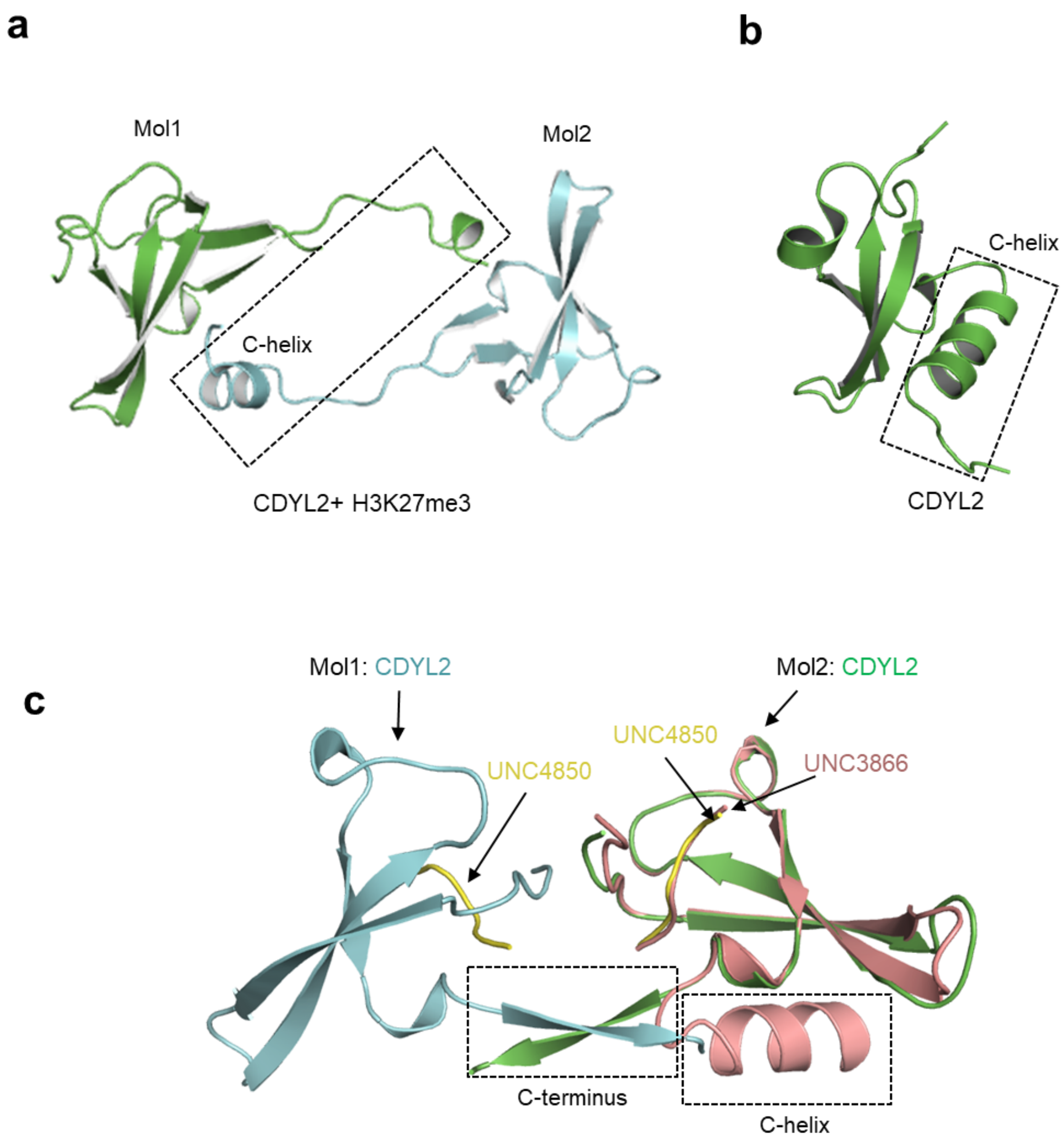


**Supplementary Figure 3 a**, Homodimer of the CDYL2 chromodomain in complex with histone H3K27me3 peptide. **b**, NMR structure of the CDYL2 chromodomain. **c**, CDYL2 forms crystal packing at the C-terminal helix with its closest neighbor (colored in cyan and green). Overlap of CDYL2-UNC3866 complex structure (colored in salmon) with one CDYL2-UNC4850 complex structure (colored in green).

a

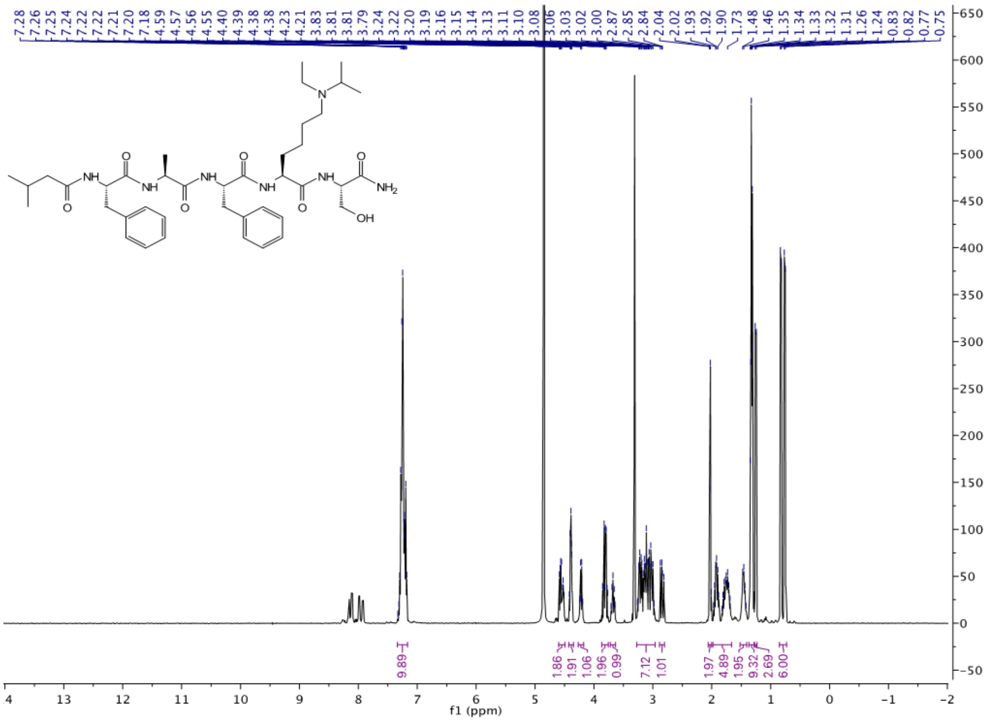


b

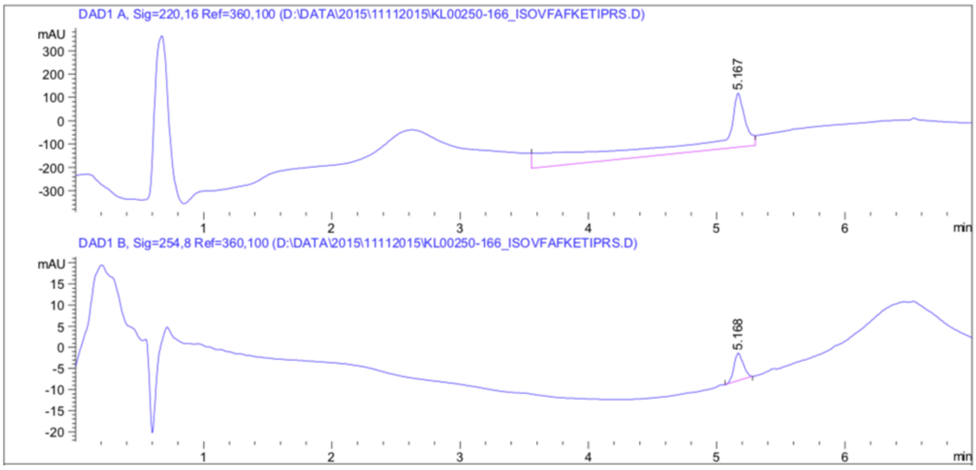


**Supplementary Figure 4** UNC4850 Characterization. **a**, $^{1}$H NMR (400 MHz, Methanol-$d_4$) δ 7.34 – 7.16 (m, 10H), 4.60 – 4.49 (m, 2H), 4.43 – 4.35 (m, 2H), 4.27 – 4.18 (m, 1H), 3.87 – 3.76 (m, 2H), 3.72 – 3.63 (m, 1H), 3.27 – 2.96 (m, 7H), 2.89 – 2.80 (m, 1H), 2.06 – 2.00 (m, 2H), 1.98 – 1.66 (m, 5H), 1.52 – 1.41 (m, 2H), 1.37 – 1.28 (m, 9H), 1.25 (d, $J$ = 7.2 Hz, 3H), 0.79 (dd, $J$ = 26.5, 6.6 Hz, 6H). **b**, MSI (ESI): 754 $[M+H]^{+}$. $t_R$ = 5.17 min.

### Supplementary Table 1 Data collection and refinement statistics

| Protein | CDY1-H3K9me3 | CDYL2-H3tK27me3 | CDYL2-H3K27me3 | CDYL2-UNC3866 | CBX7_V13A-UNC3866 | MPP8-UNC3866 | CDYL2-UNC4850 |
|---|---|---|---|---|---|---|---|
| **Diffraction data** | | | | | | | |
| Radiation source | rigaku fr-e | CLSI BL 08ID | APS BL 19ID | rigaku fr-e | rigaku fr-e | rigaku fr-e | APS BL 22ID |
| Radiation wavelength [Å] | 1.5418 | 0.9796 | 0.9792 | 1.5418 | 1.5418 | 1.5418 | 0.9790 |
| Space group | P 43 | H 3 | P 43 2 2 | P 21 21 21 | P 41 21 2 | P 21 21 21 | P 21 |
| Cell dimensions | | | | | | | |
| a, b, c [Å] | 42.45, 42.45, 37.19 | 210.69, 210.69, 67.19 | 60.79, 60.79, 178.60 | 45.90, 83.66, 115.04 | 40.19, 40.19, 83.12 | 38.69, 51.04, 73.78 | 88.84, 82.16, 128.63 |
| α, β, γ [Å] | 90, 90, 90 | 90, 90, 120 | 90, 90, 90 | 90, 90, 90 | 90, 90, 90 | 90, 90, 90 | 90, 97.44, 90 |
| Resolution limits [Å] | 23.36-1.60 (1.64-1.60) | 48.12-2.60 (2.72-2.60) | 44.65-2.70 (2.83-2.70) | 42.63-2.00 (2.05-2.00) | 22.81-1.57 (1.61-1.57) | 23.66-1.57 (1.61-1.57) | 36.25-2.70 (2.77-2.70) |
| Rmerge | 0.028 (0.334) | 0.096 (1.010) | 0.085 (1.097) | 0.124 (0.779) | 0.042 (0.508) | 0.042 (0.749) | 0.129 (0.875) |
| I/sigma | 43.6 (6.0) | 14.7 (1.9) | 26.8 (3.0) | 14.2 (2.4) | 44.2 (5.5) | 29.6 (2.7) | 9.0 (1.6) |
| Completeness [%] | 99.9 (100.0) | 100.0 (100.0) | 99.9 (99.8) | 88.8 (95.9) | 100.0 (100.0) | 99.6 (98.4) | 99.6 (98.6) |
| Redundancy | 7.0 (6.7) | 5.9 (5.9) | 13.6 (14.0) | 6.9 (6.6) | 13.1 (12.7) | 6.8 (6.6) | 3.8 (3.7) |
| **Model refinement** | | | | | | | |
| Resolution [Å] | 23.36-1.60 | 48.12-2.75 | 44.65-2.70 | 42.63-2.10 | 22.81-1.57 | 23.66-1.57 | 36.26-2.70 |
| No. Reflections used/free | 8396/411 | 27017/1877 | 9150/673 | 22151/1136 | 9344/711 | 19904/939 | 47926/2504 |
| Rwork/Rfree | 0.186/0.201 | 0.210/0.230 | 0.226/0.271 | 0.225/0.270 | 0.185/0.232 | 0.214/0.254 | 0.210/0.270 |
| No. atoms/B-factors [$Å^2$] | 659/23.9 | 3368/60.4 | 1211/89 | 3499/23.9 | 595/17.5 | 1234/24.9 | 12292/39.1 |
| Protein | 531/23.8 | 2909/61.1 | 1001/88.7 | 2976/24.1 | 467/16.1 | 1014/23.7 | 10898/39.7 |
| Ligand | 84/23.3 | 427/56.6 | 197/92.6 | 334/23.8 | 57/17.1 | 116/32.8 | 1368/35.0 |
| Water | 33/27.3 | | | 172/22.1 | 54/28.3 | 85/29.5 | |
| Others | 11/24 | 32/40.8 | 13/64.2 | 17/13.3 | 17/23.4 | 19/25.1 | 26/30.1 |
